# *PERPETUAL FLOWERING2* coordinates the vernalization response and perennial flowering in *Arabis alpina*

**DOI:** 10.1101/470567

**Authors:** Ana Lazaro, Yanhao Zhou, Miriam Giesguth, Kashif Nawaz, Sara Bergonzi, Ales Pecinka, George Coupland, Maria C. Albani

**Affiliations:** Botanical Institute, Cologne Biocenter, University of Cologne, Zülpicher Str. 47B, 50674 Cologne, Germany; Max Planck Institute for Plant Breeding Research, Carl-von-Linné-Weg 10, 50829 Cologne, Germany; Cluster of Excellence on Plant Sciences “From Complex Traits towards Synthetic Modules”, 40225 Düsseldorf, Germany; Institute of Experimental Botany, Centre of the Region Haná for Biotechnological and Agricultural Research, Šlechtitelů 31, 77900 Olomouc, Czech Republic

**Keywords:** *APETALA2*, *AP2*, juvenility, *FLOWERING LOCUS C*, *FLC*, perennial, *PERPETUAL FLOWERING 1*, *PEP1*, *PEP2*, vernalization

## Abstract

The floral repressor *APETALA2* (*AP2*) in Arabidopsis regulates flowering through the age pathway. The *AP2* orthologue in the alpine perennial *Arabis alpina*, *PERPETUAL FLOWERING 2* (*PEP2*), was previously reported to regulate flowering through the vernalization pathway by enhancing the expression of another floral repressor *PERPETUAL FLOWERING 1* (*PEP1*), the orthologue of Arabidopsis *FLOWERING LOCUS C* (*FLC*). However, *PEP2* also regulates flowering independently of *PEP1*. To characterize the function of *PEP2* we analyzed the transcriptomes of *pep2* and *pep1* mutants. The majority of differentially expressed genes were detected between *pep2* and the wild type or between *pep2* and *pep1*, highlighting the importance of the *PEP2* role that is independent of *PEP1*. Here we demonstrate that *PEP2* prevents the upregulation of the *A. alpina* floral meristem identity genes *FRUITFUL* (*AaFUL*), *LEAFY* (*AaLFY*) and *APETALA1* (*AaAP1*) which ensure floral commitment during vernalization. Young *pep2* seedlings respond to vernalization, suggesting that *PEP2* regulates the age-dependent response to vernalization independently of *PEP1*. The major role of *PEP2* through the *PEP1*-dependent pathway takes place after vernalization, when it facilitates *PEP1* activation both in the main shoot apex and in the axillary branches. These multiple roles of *PEP2* in vernalization response contribute to the *A. alpina* life-cycle.

**HIGHLIGHT:** The *Arabis alpina APETALA2* orthologue, *PERPETUAL FLOWERING2*, regulates the age-dependent response to vernalization and it is required to facilitate the activation of the *A. alpina FLOWERING LOCUS C* after vernalization.

## INTRODUCTION

Plant adaptation to environment requires the modification of developmental traits, among which flowering time is key to ensure successful production of offspring. Alpine habitats in which juvenile survival is very low are mainly dominated by perennial species (Billings and Mooney, 1968). In general, the perennial growth habit relies on the differential behavior of meristems on the same plant so that some will stay vegetative whereas others will initiate flowering (Amasino, 2009; Lazaro *et al*., 2018). The main environmental cue that promotes flowering in alpine species is the exposure to prolonged cold, a process called vernalization. Alpine environments are characterized by short growing seasons and long periods of snow coverage. Thus, to ensure reproductive success, alpine plants initiate flower buds in response to prolonged cold several months or years before anthesis (Diggle, 1997; Meloche and Diggle, 2001). However, exposure to long periods of cold does not always result in flowering. This is especially true for perennial species, as most of them have a prolonged juvenile phase and are not competent to flower at a young age (Bergonzi and Albani, 2011).

The molecular mechanisms regulating flowering in response to vernalization or to the age of the plant have been mainly studied in the annual model plant *Arabidopsis thaliana*. The MADS box transcription factor FLOWERING LOCUS C (FLC) is the major regulator of flowering in response to vernalization (Michaels and Amasino, 1999; Sheldon *et al*., 2000). FLC transcriptionally regulates floral integrator genes such as *SUPPRESSOR OF OVEREXPRESSION OF CONSTANS1* (*SOC1*), and genes involved in the age pathway, suggesting an interplay between these two pathways (Deng *et al*., 2011; Mateos *et al*., 2017). Comparative studies between Arabidopsis and the alpine perennial *Arabis alpina* demonstrated that the *FLC* orthologue in *A. alpina*, *PERPETUAL FLOWERING1* (*PEP1*), also regulates flowering in response to vernalization. In addition, *PEP1* contributes to the perennial growth habit by repressing flowering in a subset of axillary meristems after vernalization (Lazaro *et al*., 2018; Wang *et al*., 2009). Flower buds in *A. alpina* are formed during prolonged exposure to vernalizing conditions. The length of vernalization determines *PEP1* reactivation in the inflorescence. After insufficient vernalization, *PEP1* mRNA is reactivated and results in the appearance of floral reversion phenotypes such as bracts and vegetative inflorescence branches (Lazaro *et al*., 2018). In the axillary branches, the length of vernalization does not influence *PEP1* expression and *PEP1* transcript rises irrespective of the length of vernalization (Lazaro *et al*., 2018). The fate of these axillary branches is determined by a combined action of the age pathway and *PEP1* (Park *et al*., 2017; Wang *et al*., 2011).

In Arabidopsis, the age pathway is regulated by two microRNAs and their targets. MicroRNA 156 (miR156) prevents flowering at a young age and gradually decreases as the plant gets older. miR172 follows the opposite pattern and gradually accumulates during development (Wu *et al*., 2009). miR156 transcriptionally regulates a family of transcription factors named SQUAMOSA PROMOTER BINDING PROTEIN-LIKE (SPLs) (Schwab *et al*., 2005; Wu *et al*., 2009; Wu and Poethig, 2006; Xu *et al*., 2016). From these, SPL9 and SPL15 have been reported to activate the transcription of *miRNA172b*, which in turn represses the expression of a small subfamily of APETALA2-like transcription factors by a translational mechanism (Aukerman and Sakai, 2003; Chen, 2004; Hyun *et al*., 2016; Mathieu *et al*., 2009; Wu *et al*., 2009). This subfamily includes six members: AP2, TARGET OF EARLY ACTIVATION TAGGED1 to 3 (TOE1-3), SCHLAFMUTZE (SMZ), and SCHNARCHZAPFEN (SNZ) (Aukerman and Sakai, 2003; Mathieu *et al*., 2009; Schmid *et al*., 2003; Yant *et al*., 2010). *A. alpina* has a very distinct juvenile phase and the accession Pajares requires at least five weeks growth in long days before it is able to flower in response to vernalization (Bergonzi *et al*., 2013a; Wang *et al*., 2011). The role of miR156 is conserved in *A. alpina* as *miR156b* overexpressing lines block flowering in response to vernalization, while mimicry lines (MIM156), used to reduce miRNA activity, flower when vernalized at the age of three weeks (Bergonzi *et al*., 2013a). However, the complementary expression patterns of miR156 and miR172 are uncoupled in *A. alpina* (Bergonzi *et al*., 2013a). Only the accumulation of miR156 is reduced in the shoot apex as the plants get older. At this stage miR172 is not detected in the shoot apex, although plants acquire competence to flower (Bergonzi *et al*., 2013a). For flowering to occur and to observe an increase in miR172 levels in the shoot apex, exposure to vernalization is required (Bergonzi *et al*., 2013a). However, vernalization is only effective in mature plants but not in juvenile plants that express still high levels of miR156 (Bergonzi *et al*., 2013a). The initiation of flowering during cold in mature plants correlates with the gradual increase in expression of the floral organ identity genes *LEAFY* (*AaLFY*), *FRUITFUL* (*AaFUL*) and *APETALA1* (*AaAP1*) (Lazaro *et al*., 2018). In perennials, such as apple and poplar the homologues of the floral repressor *TERMINAL FLOWER1* (*TFL1*) regulate the juvenile period. Transgenic *Malus domestica* and *Populus trichocarpa* lines with reduced *TFL1* activity have a shortened juvenile phase (Kotoda *et al*., 2006; Mohamed *et al*., 2010). Similarly, the silencing of *TFL1* in *A. alpina* allows flowering in young vernalized seedlings (Wang *et al*., 2011). Interestingly, these lines can flower after being vernalized for a short time (6 instead of 12 weeks). These results suggest again an interplay between the age and the vernalization pathways.

In Arabidopsis, *AP2* influences a variety of developmental processes, including flowering time through the age pathway and floral development (Yant *et al*., 2010). Strong *AP2* mutant alleles, such as *ap2-12*, flower early in both long days and short days (Yant *et al*., 2010). Similarly, the *A. alpina* orthologue of *AP2*, *PEP2* has been reported to have a flowering time phenotype (Bergonzi *et al*., 2013a). *pep2* mutants flower without vernalization and show compromised perennial traits, similar to *pep1-1* mutant plants (Bergonzi *et al*., 2013a; Wang *et al*., 2009). The effect of *PEP2* on flowering was first related to the vernalization pathway as it promotes the expression of *PEP1* (Bergonzi *et al*., 2013a). In 2-week-old *pep2-1* seedlings, *PEP1* transcript levels are reduced compared to wild type plants (Bergonzi *et al*., 2013a). However, *PEP2* also has a *PEP1*-independent role in the regulation of flowering time in *A. alpina* as flowering is accelerated in the *pep1-1 pep2-1* double mutant compared to the single mutants (Bergonzi *et al*., 2013a).

Here, we show that during vernalization *PEP2* represses the expression of the floral meristem identity genes *AaFUL*, *AaLFY* and *AaAP1*. Vernalization accelerates flowering in young *pep2-1* plants, indicating that *PEP2* regulates the age-dependent response to vernalization. In addition, we report that the *PEP1*-dependent role of *PEP2* takes place after vernalization because *PEP2* is required to activate *PEP1* after the return to warm temperatures. The involvement of *PEP2* in two different aspects of the vernalization response contribute to the perennial life-cycle of *A. alpina*.

## RESULTS

### *PEP2* influences the expression of genes involved in many plant physiological and developmental responses including flowering

To provide an overview of the role of *PEP2* in *A. alpina* we performed an RNAseq analysis. We compared the transcriptomes of apices of 3-week-old *pep2-1* and *pep1-1* mutants to the wild type (Pajares). Three-week old wild type and mutant plants are vegetative and have not undergone the transition to flowering (Bergonzi *et al*., 2013a; Lazaro *et al*., 2018; Park *et al*., 2017; Wang *et al*., 2011). Among transcriptomes, the majority of differentially expressed genes were detected in *pep2-1*. A total of 253 genes were up-regulated and 223 genes were down-regulated in *pep2-1* compared to the wild type (Fig. 1A, B; Dataset 1). In contrast, only 47 genes were up-regulated and 98 genes were down-regulated in *pep1-1* compared to the wild type (Fig. 1A, B; Dataset 2). The genes differentially expressed between *pep1-1* and the wild type are influenced by *PEP1*, whereas the ones differentially expressed between *pep2-1* and the wild type are affected by *PEP2* both through the *PEP1*-dependent and *PEP1*-independent pathway. To identify genes influenced by *PEP2* through the *PEP1*-independent pathway we compared the transcriptomes of *pep2-1 vs pep1-1* (Fig. 1C, D; Dataset 3). A total of 504 genes were significantly up- and 251 genes significantly down-regulated in *pep2-1* compared to *pep1-1* (Fig. 1C, D). Interestingly, the number of differentially expressed genes detected between *pep2-1* and *pep1-1* was higher than the ones detected when single mutants were compared to the wild type. Gene Ontology (GO) analysis demonstrated that the most enriched category for the up regulated genes in *pep2-1* compared to the wild type and in *pep2-1* compared to *pep1-1* was the biosynthesis of glucosinolates, which are involved in defense against herbivore attack and pathogens (Fig. S1) (Keith and Mitchell-Olds, 2017). The overlap in overrepresented GO categories in the set of genes up-regulated in *pep2-1* in comparison to either the wild type or the *pep1-1* mutant was very high, which is to be expected as more genes were up-regulated in *pep2-1* compared to the wild type than in *pep1-1* compared to the wild type (Fig. 1A and Fig. S1A, B). Among the commonly enriched categories for down-regulated genes in *pep2-1*, we found apoptosis and protein desumoylation (Fig. S1C, D).

**Fig. 1.**
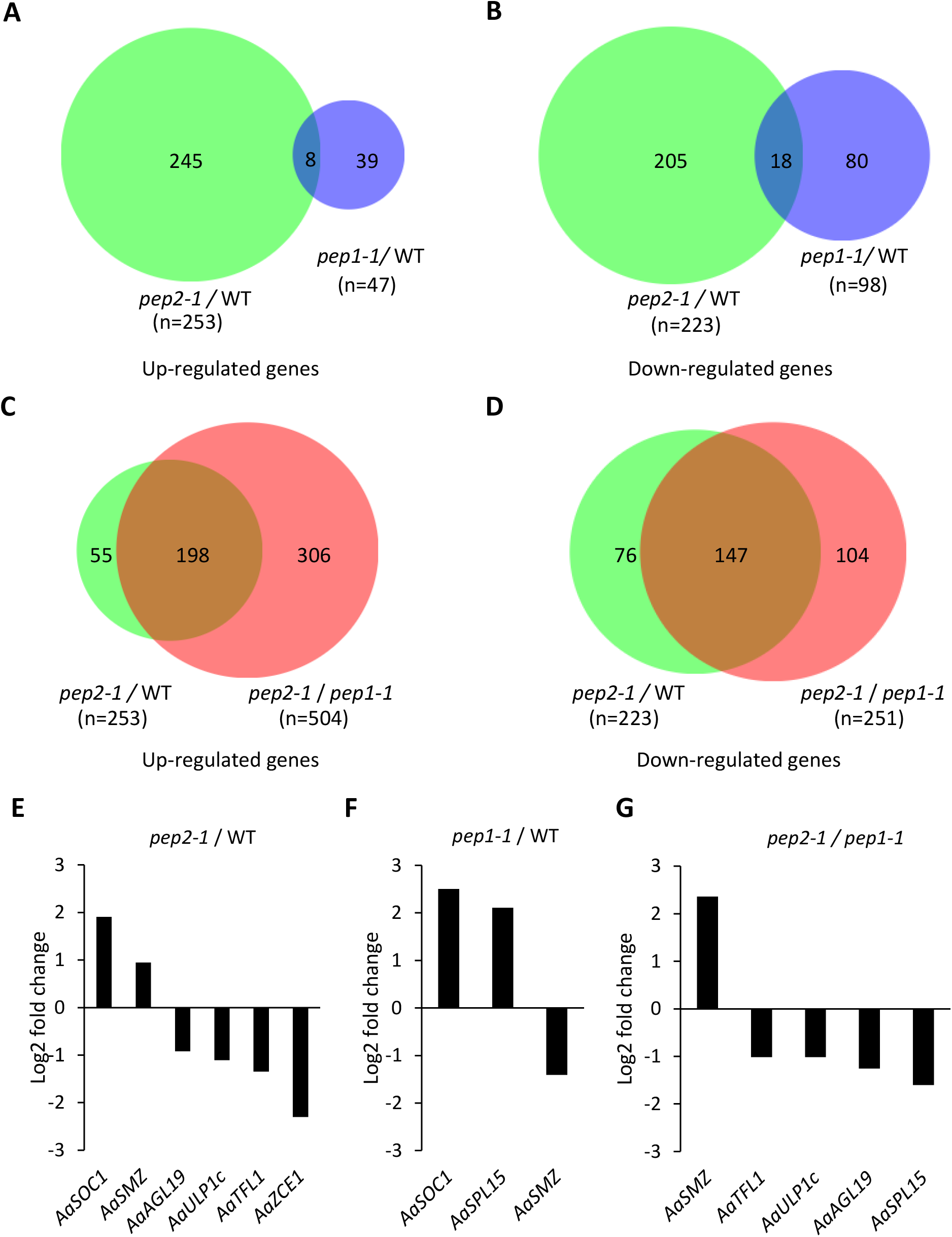
Differentially expressed genes in *pep2* and *pep1 A. alpina* mutants. **(A and B)** Venn diagram of significantly up-regulated (A) and down-regulated (B) genes in *pep2-1* compared to the wild type (WT) and *pep1-1* compared to the WT. **(C and D)** Venn diagram of significantly up-regulated (C) and down-regulated (D) genes in *pep2-1* compared to the WT and in *pep2-1* compared to *pep1-1*. **(E-G)** Flowering time genes differentially expressed in *pep2-1* compared to the WT (E), *pep1-1* compared to the WT (F) and *pep2-1* compared to *pep1-1* (G). Expression values are based on RNA-sequencing.

Floral activators and repressors were identified among the differentially expressed genes in *pep2-1*. For example, the *A. alpina* orthologue of *SOC1* (*AaSOC1*) was up-regulated in *pep2-1* compared to the wild type (Fig. 1E and Dataset 1). This effect of *PEP2* on *AaSOC1* is through *PEP1* as *AaSOC1* was differentially expressed between *pep1-1* and the wild type, but not in *pep2-1 vs pep1-1* (Fig. 1F, G and Dataset 2 and 3; Mateos et al, 2017). The regulation of *AaSMZ* by *PEP2* is different than by *PEP1*. *AaSMZ* was up-regulated in *pep2-1* compared to the wild type and down-regulated in *pep1-1* compared to the wild type (Fig. 1E, F and Dataset 1 and 2). In contrast, *AaSPL15* was up-regulated in the *pep1-1* mutant compared to the wild type and not in *pep2-1* compared to the wild type, indicating that *PEP2* does not control *AaSPL15* expression (Fig. 1E, F and Dataset 1 and 2). Among the flowering time genes involved in the *PEP1*-independent role of *PEP2* were the floral repressor *AaTFL1* and *AGAMOUS-LIKE 19* (*AaAGL19*). *AaTFL1* was down-regulated when we compared *pep2-1* to both the wild type and *pep1-1*, suggesting that the effect of *PEP2* on *AaTFL1* is independent of *PEP1* (Fig. 1E-G and Dataset 1-3). Similarly, *AGAMOUS-LIKE 19* (*AaAGL19*) transcripts were down-regulated specifically in the *pep2-1* mutant (Fig. 1E-G and Dataset 1-3). We also found the SUMO protease *AaULP1c* and the orthologue of *CIS-CINNAMIC ACID-ENHANCED 1* (*AaZCE1*) being differentially expressed specifically in *pep2-1* (Fig. 1C-E and Dataset 1 and 2). Interestingly, both *ULP1c* and *ZCE1* in Arabidopsis regulate flowering through *FLC*. Mutations in this desumoylating enzyme *ULP1c* and its homolog, *ULP1d*, show an early flowering phenotype in Arabidopsis that can at least partially be due to *FLC* down regulation (Castro *et al*., 2016; Conti *et al*., 2008). *ZCE1* is involved in the regulation of plant growth and development by *cis*-phenylpropanoids and it has been shown to regulate bolting time by enhancing *FLC* expression (Guo *et al*., 2011).

### *PEP2* can complement the Arabidopsis *ap2* mutant

Both the *pep2* mutant in *A. alpina* and the *ap2* mutant in Arabidopsis show early flowering and similar floral defects, including the absence of petals and the transformation of sepals to carpels (Bergonzi *et al*., 2013b; Bowman *et al*., 1991; Nördstrom *et al*., 2013). To check if both genes have common functions, we expressed *PEP2* in the *ap2-7* mutant background under the control of its own promoter. We fused a 7.4 Kb *PEP2* genomic region spanning 4 Kb upstream of the translational start and 1.2 Kb downstream of the translational stop to the VENUS fluorescent protein, at the N- or C-terminus. Transgenic lines were first obtained in Col background. Homozygous lines obtained for the N-terminal VENUS (Col *ProPEP2::VENUS::PEP2* N6-1-3) and the C-terminal VENUS (Col *ProPEP2::PEP2::VENUS* C2-1-9) were subsequently crossed to *ap2-7*. When grown in SDs, the *PEP2* constructs complemented the early flowering phenotype of the *ap2-7* mutant (Fig. 2A-D). Moreover, the homeotic defects of the *ap2* mutant were restored by *PEP2*, indicating that the *A. alpina PEP2* gene regulates in a similar way to *AP2* flowering time and floral organ identity (Fig. 2E-H).

**Fig. 2.**
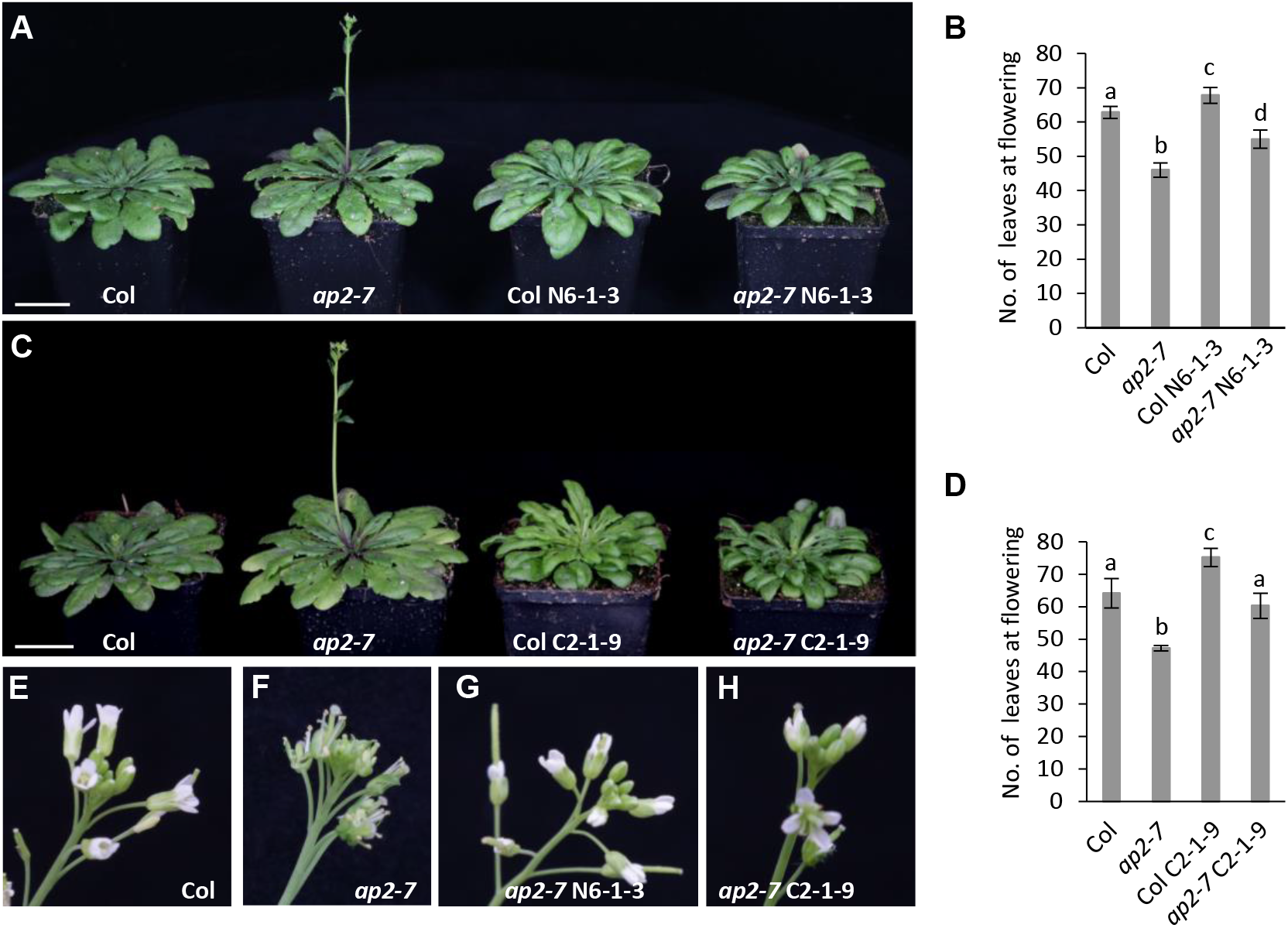
*PEP2* can complement the flowering and floral phenotype of the Arabidopsis *ap2-7* mutant. **(A and B)** Phenotypes of Col wild type, the *ap2-7* mutant, the Col *ProPEP2::VENUS::PEP2* N6-1-3 and the *ap2-7 ProPEP2::VENUS::PEP2* N6-1-3 lines grown in SDs (A) and number of leaves at flowering (B). **(C and D)** Col, the *ap2-7* mutant, the Col *ProPEP2::PEP2::VENUS* C2-1-9 and the *ap2-7 ProPEP2::PEP2::VENUS* C2-1-9 lines grown in SDs (C) and number of leaves at flowering (D). (A and C) Whole plant pictures were taken 57DAG. Bar = 3cm. In B and D asterisks stand for significant differences determined by a Student T-test (p-value<0.01). Error bars indicate s.d.m. **(E to F)** Inflorescence of Col wild type (E) *ap2-7* (F), *ap2-7 ProPEP2::VENUS::PEP2* N6-1-3 (G) and *ap2-7 ProPEP2::PEP2::VENUS* C2-1-9 (H) taken 73DAG in SDs.

To test whether the effect of *PEP2* on *PEP1* expression was conserved in Arabidopsis for *AP2* and *FLC*, we combined the *ap2-7* mutation with the strong *FRI* allele from the San Feliu-2 (Sf-2) accession, which enhances Col *FLC* expression. Although the *ap2* mutation reduced the number of leaves to half in the *FRI* Sf-2 background, the expression of *FLC* is not altered in the apices of these plants at different developmental stages (before, during, or after 40 days of vernalization) (Fig. S2). These results indicate that, although the role of *AP2* and *PEP2* regarding flowering time regulation and floral organ identity is conserved, *AP2* does not regulate *FLC* expression in a *FRI* Sf-2 background (Fig. S2B).

### *PEP2* regulates the age-dependent response of *A. alpina* to vernalization

We then investigated whether the *PEP1*-independent role of *PEP2* was similar to the one of *AP2* in Arabidopsis and therefore whether it regulated flowering through the age pathway. We first analyzed the accumulation of miR156 and the transcript level of the *A. alpina SPL5, 9* and *15* (*AaSPL5, 9* and *15*) in the apices of *pep2-1* and wild type seedlings grown for 3, 4, and 6 weeks in LDs (Fig. S3). miR156 accumulation in the shoot apex was downregulated in older seedlings but a similar pattern was observed in *pep2-1* and the wild type (Fig. S3A). Transcript levels of *AaSPL5, 9* and *15* were upregulated in older plants (Fig. S3B-D). For *AaSPL5* and *15* we observed no significant differences between *pep2-1* and the wild type, whereas *AaSPL9* mRNA levels differed between the two genotypes only in 6-week-old seedlings (Fig. S3B-D). These results are consistent with previous studies in Arabidopsis demonstrating that *AP2* regulates flowering through the age-pathway downstream of miR156 and the *SPL*s. As it was previously shown that the age-dependent effect on flowering in *A. alpina* is only apparent after vernalization (Bergonzi *et al*., 2013a; Wang *et al*., 2011), we tested whether *PEP2* has an age-dependent role in vernalized plants. For this we vernalized 3-week-old wild type and *pep2-1* seedlings for 12 weeks and measured flowering time after the return to warm temperatures. We also included the *pep1-1* mutant in this experiment to rule out a *PEP1*-dependent effect of *PEP2* on flowering time. In accordance to previous studies, under these conditions the wild type did not flower after vernalization and only grew vegetatively (Fig. 3; and Wang et al., 2011; Bergonzi et al, 2013). Interestingly, *pep2-1* flowered with an average of 18 leaves and 17 days after vernalization, whereas vernalized *pep1-1* flowered with 27 leaves similar to non-vernalized *pep1-1* plants grown continuously in long days (Fig. 3 and Fig. S4; Wang et al., 2009; Bergonzi et al., 2013). This data suggests that vernalization accelerates flowering in young *pep2-1* but not in *pep1-1* plants. The flowering time phenotype of the mutants is also in contrast to one in long days where *pep1-1* flowers earlier than *pep2-1* (Bergonzi et al., 2013). Overall these results suggest that *PEP2* regulates the age-dependent response to vernalization in a *PEP1*-independent manner.

**Fig. 3.**
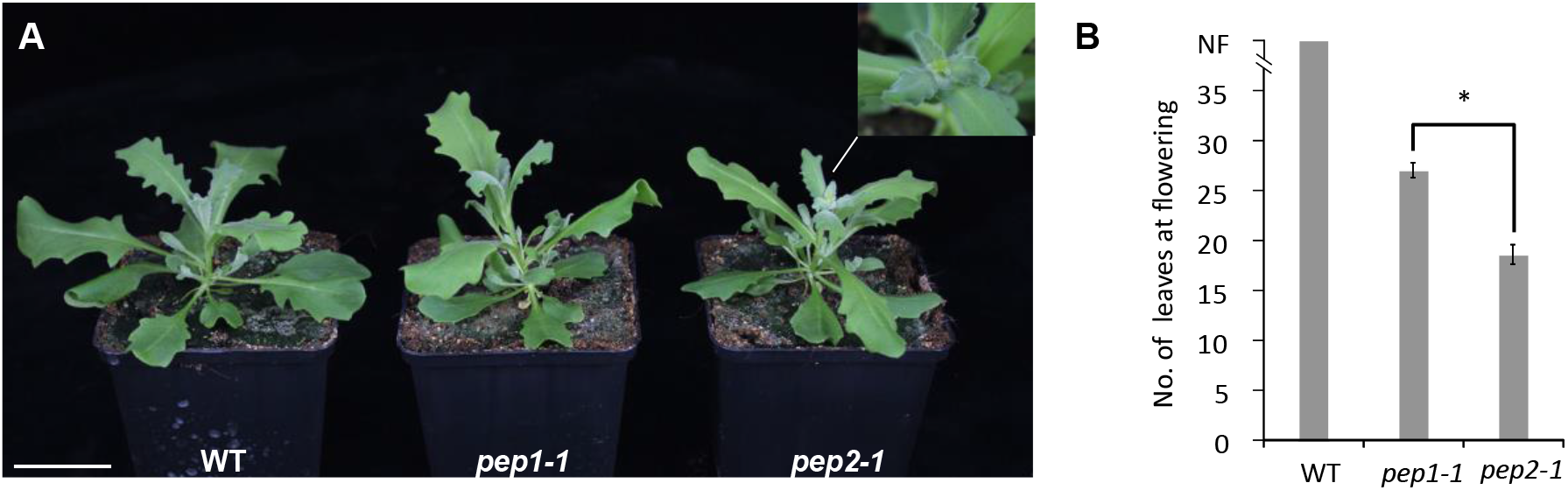
*PEP2* regulates the age-dependent response of *A. alpina* to vernalization. **(A)** Picture of 3-week-old wild type (WT), *pep1-1* and *pep2-1* vernalized for 12 weeks followed by 2 weeks in LDs. Bar = 5cm. **(B)** Flowering time demonstrated as the number of leaves at flowering of 3-week-old WT, *pep1-1* and *pep2-1* mutants vernalized for 12 weeks. The WT did not flower (NF). The asterisk stands for a significant difference in the total leaf number determined by a Student T-test (p-value<0.01). Error bars indicate s.d.m.

To understand how the young *pep2-1* plants accelerate flowering in response to vernalization, we analyzed the expression of *PEP1*, *AaSOC1*, *AaFUL*, *AaTFL1*, *AaLFY* and *AaAP1*. Three-week-old wild type, *pep2-1* and *pep1-1* apices from the main shoot were harvested before and during vernalization at 4, 8 and 12 weeks. In accordance with previous results obtained in seedlings, unvernalized 3-week-old *pep2-1* plants showed lower *PEP1* mRNA levels than the wild type (Fig. 4A; Bergonzi et al., 2013). Nevertheless, *PEP1* transcript level was influenced in a similar way in *pep2-1* and wild type plants and *PEP1* was silenced after four weeks in cold (Fig. 4A). This data suggests that, despite the initial difference in *PEP1* expression, the lack of *PEP2* does not influence *PEP1* transcription in young apices during vernalization. The expression of *AaSOC1* was gradually up-regulated during vernalization, following the same pattern in the three genotypes (Fig. 4B). In contrast, *AaFUL*, *AaLFY* and *AaAP1* showed a differential increase in the wild type and the mutants after 8 and 12 weeks in vernalization (Fig. 4C, E and F). In young wild type plants *AaFUL*, *AaLFY* and *AaAP1* mRNA levels did not rise, indicating that flowering had not been initiated (Fig. 4C, E and F). Moreover, the *pep2-1* mutant showed higher levels of *AaLFY* and *AaAP1* than *pep1-1* after 12 weeks in vernalization (Fig. 3 and 4E and F). Interestingly, the *pep2-1* mutant also showed reduced expression of *AaTFL1* at the end of the cold treatment compared to *pep1-1* (Fig. 4D). Taken together, our results indicate that *PEP2* activates *AaTFL1* and represses *AaFUL*, *AaLFY* and *AaAP1* in young apices during vernalization (Fig. 3). This role of *PEP2* is independent of *PEP1*, given that *pep1-1* plants vernalized at a young age flowered later than *pep2-1* and that *PEP1* expression was reduced to the same extent in wild type and *pep2-1* during vernalization (Fig. 3 and 4A).

**Fig. 4.**
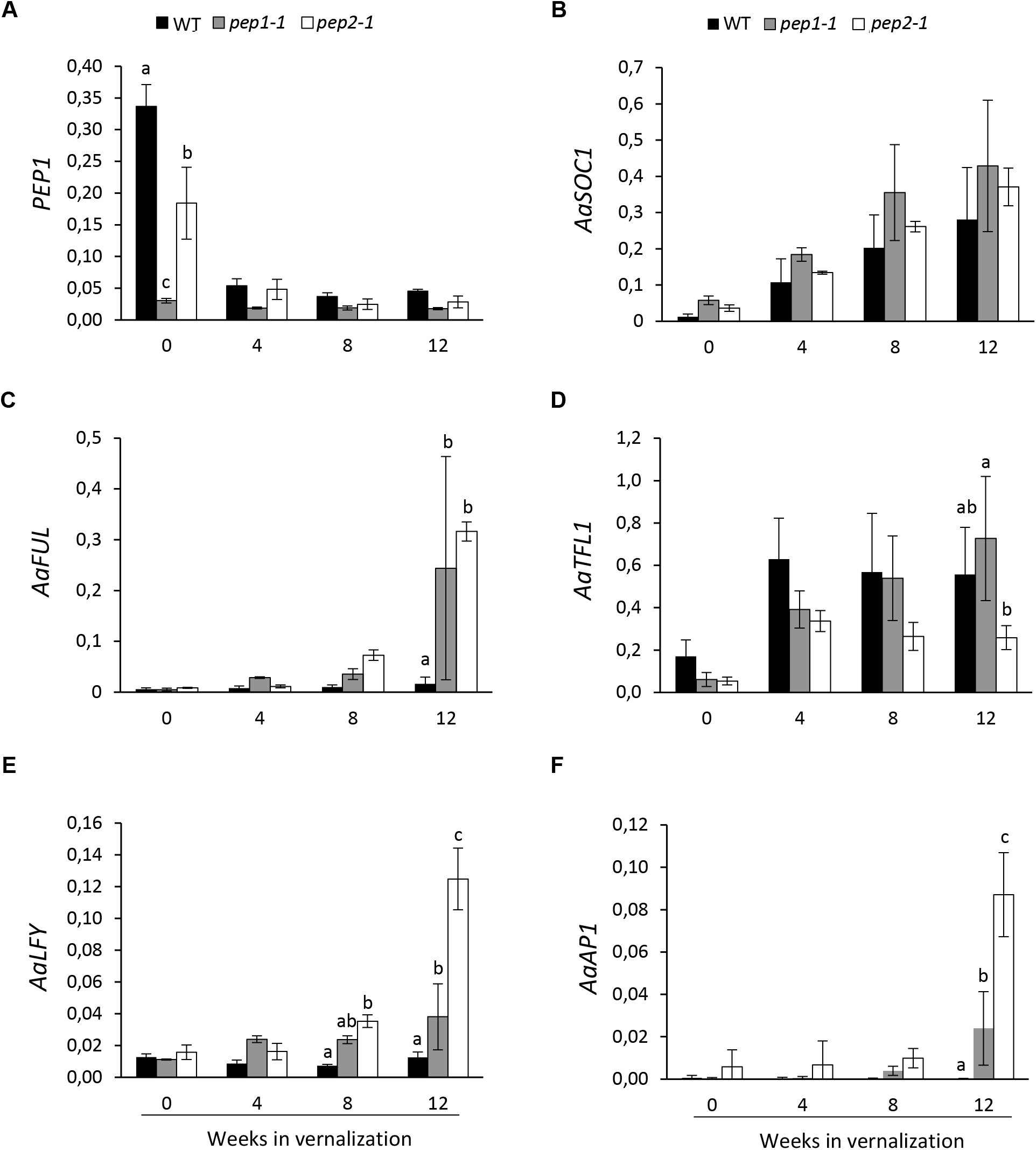
*PEP2* regulates *AaFUL*, *AaTFL1*, *AaLFY* and *AaAP1* expression during vernalization. Relative expression of *PEP1* **(A)**, *AaSOC1* **(B)**, *AaFUL* **(C)**, *AaTFL1* **(D)**, *AaLFY* **(E)** and *AaAP1* **(F)**. Three-week-old wild type (WT), *pep1-1* and *pep2-1* shoot apices were harvested before and during 12 weeks of vernalization. Letters stand for significant differences between WT, *pep1- 1* and *pep2-1* at each time point determined by multiple pairwise comparisons using Benjamini-Hochberg-corrected p-values (α-value of 0.05). Graphs with no letters show no significant differences. Error bars indicate s.d.m.

To investigate whether the transcriptional regulation of *PEP2* on these floral meristem identity genes was also conserved in adult plants, we tested the expression of *AaFUL*, *AaLFY* and *AaAP1* during vernalization. Six-week-old wild type and *pep2-1* plants were exposed to 12 weeks of cold and the mRNA levels of *AaLFY*, *AaAP1* and *AaFUL* was analyzed in the shoot apex before vernalization and 1, 3, 5, 8 and 12 weeks into vernalization. *AaFUL* mRNA levels were higher in *pep2-1* than in the wild type already after 8 weeks in vernalization (Fig. S4A). For *AaLFY* and *AaAP1* expression a significant increase was observed in the *pep2-1* mutant only at the end of the 12 weeks of cold (Fig. S4B and C). Overall these results suggest that *PEP2* delays flowering by keeping *AaFUL*, *AaLFY* and *AaAP1* repressed at the end of the 12 weeks of vernalization, when *PEP1* has already been silenced in the apices of both young and adult plants.

### *PEP2* is required to activate *PEP1* expression after vernalization

To test the *PEP1*-dependent role of *PEP2* we exposed the *pep2-1* mutant and the wild type plants to different lengths of vernalization. Both genotypes were grown for 5 weeks in LDs, vernalized for 8, 12, 18 and 21 weeks, and transferred back to LD glasshouse conditions (Fig. 5A and B). The *pep2-1* mutant showed a reduction in the number of days to flower emergence compared to the wild type in all durations of cold (Fig. 5C). In addition, inflorescences in *pep2-1* showed reduced floral reversion phenotypes and enhanced commitment of inflorescence branches to flowering (Fig. 5D-G). These results indicate that *PEP2* regulates flowering time and inflorescence architecture in *A. alpina*. However, the response of *pep2-1* still varied with the length of vernalization suggesting that other floral repressors might contribute to flowering in response to vernalization. Also, *PEP2* is required to maintain axillary shoots that are located just below the inflorescence in a vegetative state as all axillary branches in the *pep2-1* mutant commit to reproductive development (Fig. 5; Bergonzi et al., 2013).

**Fig. 5.**
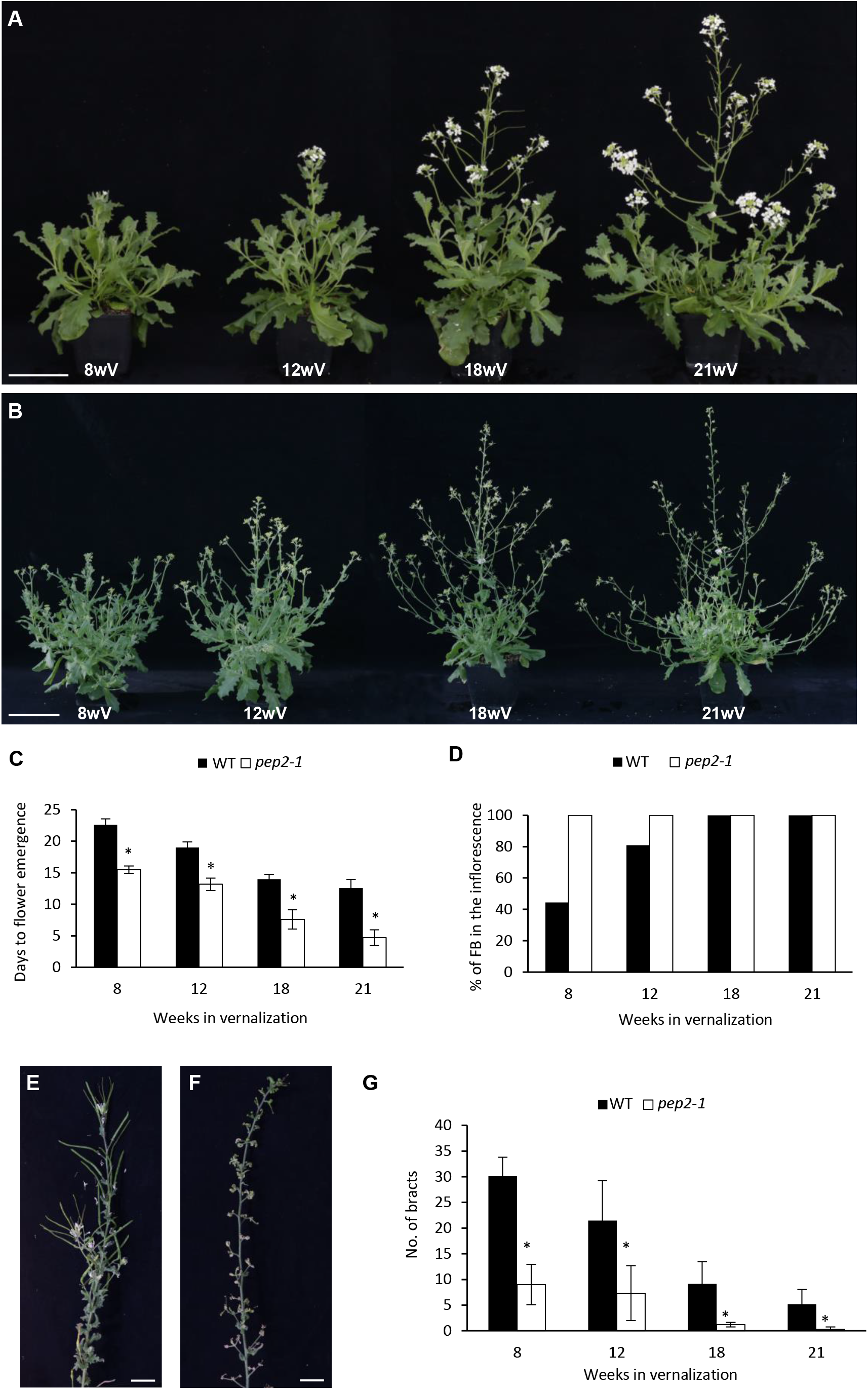
The *pep2* mutant plants flower earlier than the wild type and show reduced reverted phenotypes. **(A)** Wild type (WT) plants exposed to several durations of vernalization (8, 12, 18 and 21 weeks) followed by 3 weeks in LDs. **(B)** *pep2-1* mutant plants exposed to several durations of vernalization (8, 12, 18 and 21 weeks) followed by 3 weeks in LDs. Bar = 10cm. **(C)** Time to flower emergence of WT and *pep2-1* plants exposed to different durations of vernalization measured as the number of days to the first open flower. **(D)** Percentage of flowering inflorescence branches (FB) in the WT and the *pep2-1* mutant exposed to 8, 12, 18, and 21 weeks of vernalization at the time the last flower in the inflorescence opened. **(E)** WT reverted inflorescence in plants vernalized for 8 weeks. **(F)** *pep2-1* mutant inflorescence in plants vernalized for 8 weeks. Bar = 2cm. **(G)** Number of bracts within the inflorescence of the WT and the *pep2-1* mutant exposed to 8, 12,18 and 21 weeks of vernalization at the time the last flower in the inflorescence opened. This experiment was performed together with the *pep1-1* mutant in an experiment previously published (Figure 6 in Lazaro et al., 2018). Data for the WT control is similar between the two papers. Asterisks stand for significant differences between the wild type and the *pep2-1* mutant at each time point determined by multiple pairwise Bonferroni tests (α-value of 0.05). Error bars indicate s.d.m.

As shown previously in the wild type, *PEP1* mRNA is up-regulated in the shoot apical meristem of the main shoot after a non-saturating vernalization (Fig. 6; Wang et al., 2009; Lazaro et al., 2018). This unstable silencing of *PEP1* mRNA after cold was abolished in the *pep2-1* mutant, suggesting that *PEP2* is required to activate *PEP1* expression in the shoot apical meristem after insufficient vernalization (Fig. 6). The role of *PEP2* in the activation of *PEP1* after vernalization is also observed in the axillary branches. All axillary branches in the *pep2-1* mutant committed to flowering (Fig. 5B) and showed very low expression of *PEP1* when compared to wild type vegetative branches (Fig. 6). These results suggest that the major contribution of *PEP2* is to activate *PEP1* transcription after vernalization, both in the shoot apical meristem and in the vegetative axillary branches.

**Fig. 6.**
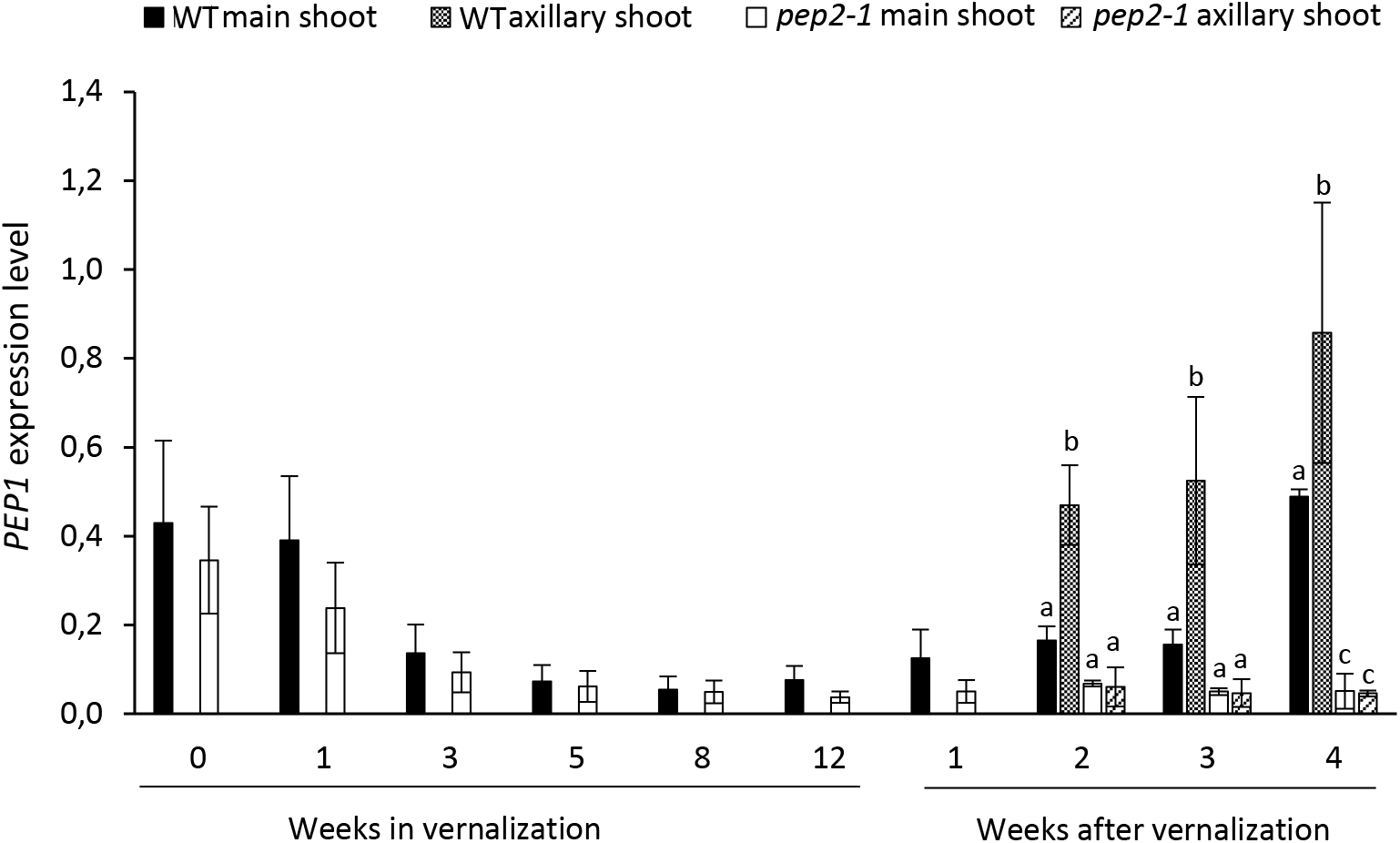
*PEP2* is required to activate *PEP1* expression after vernalization. Relative expression of *PEP1* in the shoot apical meristem and in the vegetative axillary meristems of the wild type (WT) and the *pep2-1* mutant before, during and after 12 weeks of vernalization. Asterisks stand for significant differences between the WT and *pep2-1* at each time point determined by multiple pairwise comparisons using Benjamini-Hochberg-corrected p-values (α-value of 0.05). Detailed information on significant differences can be found in Table S3. Error bars indicate s.d.m.

## MATERIALS AND METHODS

### Plant material, growth conditions and phenotyping

The *A. alpina* genotypes used in this paper were Pajares (wild type), the *pep2-1* mutant and the *pep1-1* mutant. The accession Pajares was collected in the Cordillera Cantábrica mountains in Spain at 1,400 meters altitude (42°59′32″ N, 5°45′32″ W). Both the *pep2-1* and the *pep1-1* mutant were isolated from an EMS mutagenesis in the Pajares background (Bergonzi *et al*., 2013a; Nordstrom *et al*., 2013; Wang *et al*., 2009). For the phenotypic analysis plants were grown in LDs (16 h light and 8 h dark) under temperatures ranging from 20°C during the day to 18°C during the night. All vernalization treatments were performed at 4°C in SD conditions (8 h light and 16 h dark).

Flowering time in the young wild type, *pep2-1* and *pep1-1* plants was scored as the number of leaves at flowering and as the number of days to the first open flower after vernalization. Plants were grown for 3 weeks in LD cabinets, vernalized for 12 weeks, and moved back to LDs after cold.

The characterization of flowering time and inflorescence traits with different vernalization durations in the *pep2-1* mutant was performed together with the wild type and the *pep1-1* mutant in an experiment previously published (Fig. 6 in Lazaro et al., 2018). The same data for control wild type plants was used in Lazaro et al., 2018. Plants were grown for five weeks in LD greenhouse, vernalized for 8, 12, 18 and 21 weeks, and moved back to LD greenhouse conditions on the same day. Flowering time was measured by recording the date on which the first flower opened after vernalization. The number of flowering and vegetative branches and the number of bracts in the inflorescence was measured at the end of flowering except for plants vernalized for 8 weeks when the measurements took place 14 weeks after vernalization.

The Arabidopsis genotypes used in this paper were Columbia-0 (Col-0) wild type, *ap2-7* and Col *FRI* San Feliu-2 (Sf-2) (Lee and Amasino, 1995). The *ap2-7* mutant was crossed to the Col *FRI* Sf-2 and the *FRI ap2-7* plants were isolated from a selfed F2 progeny that showed *ap2* homeotic defects and late flowering.

For the flowering time experiments in Arabidopsis the total leaf number (rosette and cauline leaves) was scored at the time the first flower opened.

### Construction of plasmids and plant transformation

To obtain the *ap2-7 PEP2*–VENUS transgenic plant, a 7.4 Kb *PEP2* genomic region spanning 4 Kb upstream of the translational start and 1.195 bp downstream of the translational stop was cloned by PCR (NCBI accession number LT669794.1). Subsequently, the VENUS:9Ala coding sequence was inserted either after the ATG or before the STOP codon of *PEP2* by employing the polymerase incomplete primer extension (PIPE) method (Klock *et al*., 2008). Primers used for PIPE-cloning are summarized in Table S1. The generated recombinant DNA fragments were integrated in the pEarlyGate301 binary vector and transformed into Col through Agrobacterium mediated floral dip (Clough and Bent, 1998). Selected homozygous lines, Col *ProPEP2::VENUS::PEP2* N6-1-3 and Col *ProPEP2::PEP2::VENUS* C2-1-9, were crossed to *ap2-7*.

### Gene Expression Analysis

Gene expression analysis was performed on the wild type, *pep1-1* and *pep2-1*. For *pep2-1* samples, homozygous plants were selected after genotyping from a segregating population using a CAP marker (Primer F: CAGCTGCACGGTATGTTTTTC, primer R: GCTTTGTCATAAGCCCTGTG, and NdeI digestion).

For the analysis of the *PEP1* expression pattern, the wild type and *pep2-1* were grown for 6 weeks in LDs and vernalized for 12 weeks. Main shoot apices were harvested before vernalization, during vernalization, and after vernalization (1, 2, 3 and 4 weeks after the plants returned to warm temperatures). Axillary vegetative apices were harvested from plants growing in LDs 2, 3 and 4 weeks after vernalization. An average of 10 apices were pooled in each sample.

The expression of *PEP1*, *AaSOC1, AaFUL*, *AaTFL1*, *AaLFY* and *AaAP1* transcripts in the young and adult wild type, *pep1-1* and *pep2-1* was detected in seedlings grown for 3 (young) or 6 weeks (adult) in LDs and vernalized for 12 weeks. Main shoot apices were harvested before vernalization and during cold, at 4, 8 and 12 weeks in vernalization. For the analysis of *AaSPL5*, *AaSPL9* and *AaSPL15* and miR156, the main shoot apex was harvested from 3-, 4- and 6-week old wild type and *pep2-1* plants growing in LDs. An average of 14 apices were pooled in each sample. Expression levels were normalized to both *AaPP2A* and *AaRAN3*, except for miR156 which was normalized to snoR101.

The expression of *FLC* transcript was analyzed in the shoot apex of *FRI* and *FRI ap2-7* plants grown for 10 days before vernalization, during 40 days of vernalization and 10 and 20 days after the return to LD glasshouse conditions. Expression levels were normalized to *UBC21*. Total plant RNA was extracted using the RNeasy Plant Mini Kit (Qiagen), and a DNase treatment was performed with Ambion DNA-free kit (Invitrogen) to reduce any DNA contamination. Total RNA (1.5 µg) was used to synthesize cDNA through reverse transcription with SuperScript II Reverse Transcriptase (Invitrogen) and oligo dT(18) as primer. 2 µl of a cDNA dilution (1:5) was used as the template for each quantitative PCR (qPCR). For the analysis of miR156 and the SPLs, total RNA was extracted using the miRNeasy^®^ Mini Kit (Qiagen), and a DNAse treatment was performed with Ambion DNA-Free kit (Invitrogen) to reduce DNA contamination. 200 ng of RNA was used for reverse transcription of miR156 and SnoR101 using miR156 and snoR101 specific primers. qPCRs were performed using a CFX96 and CFX384 Real-Time System (Bio-Rad) and the iQ SYBR Green Supermix detection system. Each data point was derived from 2 or 3 independent biological replicates and is shown as mean ± s.d.m.

Primers used for qPCR for *PEP1*, *AaSOC1*, *AaFUL*, *AaTFL1*, *AaLFY*, *AaAP1*, *AaSPL5*, *AaSPL9*, *AaSPL15*, *AaPP2A, AaRAN3*, miR156 and SnoR101 were described previously (Bergonzi *et al*., 2013a; Lazaro *et al*., 2018; Wang *et al*., 2011; Wang *et al*., 2009; Mateos *et al*., 2017). Primers used for qPCR for *FLC* and *UBC21* were also described elsewhere (Crevillen *et al*., 2013; Czechowski *et al*., 2005).

### Statistical analysis

Statistical analyses were performed using the R software. To detect significant differences in gene expression we controlled for a false discovery rate of 0.05 when conducting multiple pairwise comparisons by using Benjamini-Hochberg-corrected p-values. Treatments with significant differences are depicted with letters or asterisks. For the *pep2-1* physiological analysis we conducted multiple pairwise Bonferroni tests (α = 0.05) to detect significant differences between the wild type and *pep2-1*. Here, a nonparametric test could not be conducted due to ties created during rank assignment.

### RNAseq analysis

For differential gene expression analysis, we used the RNA sequencing method on apices from the 3-week-old wild type, the *pep2-1* and the *pep1-1* mutant. *pep2-1* homozygous plants were genotyped from a segregating population using the CAP marker described above. RNA was isolated as described above and total RNA integrity was confirmed on the Agilent BioAnalyzer. The library preparation and sequencing were performed at the Max Planck Genome Center Cologne, Germany (https://mpgc.mpipz.mpg.de/home/). RNA sequencing was performed with three biological replicates per sample. The libraries were prepared from 1 mg total RNA using the TruSeq RNA kit (Illumina) and sequenced 100-bp single-end reads on HiSeq2500 (Illumina). Reads from all samples were mapped on *A. alpina* reference genome (Willing *et al*., 2015) using TopHat (Trapnell *et al*., 2009) with default parameters. Afterwards, CuffDiff (Trapnell *et al*., 2010) was used to estimate the mRNA level of each gene by calculating fragments per kilobase of exon model per million reads mapped (FPKM). To calculate the differential gene expression among the samples FPKM values were used. A log2 fold change (L2FC) ≥ 1 for up-regulated genes and L2FC ≤ −1 for down-regulated genes, both with q-value (adjusted p-value) ≤ 0.05 was used for further analysis.

GO enrichment was performed with the BiNGO plug-in (Maere *et al*., 2005) implemented in Cytoscape V3.5.1 (Cline *et al*., 2007). A hypergeometric test was applied to determine the enriched genes and the Benjamini–Hochberg FDR correction (Benjamini and Hochberg, 1995) was performed in order to limit the number of false positives. The FDR was set up to 0.05.

Sequencing data from this study have been deposited in Gene Expression Omnibus (GEO) under accession number GSE117977. Sequences of genes studied can be found in the GenBank/EMBL databases under the following accession numbers: *PEP2* (AALP_AA7G245300), cDNA of *PEP1* (FJ755930), coding sequence of *AaLFY* (JF436956), coding sequence of *AaSOC1* (JF436957), *AaAP1* (AALP_AA2G117200), coding sequence of *AaTFL1* (JF436953), *AaFUL* (Aa_G837900).

## DISCUSSION

Understanding the role of prolonged exposure to low temperatures in flowering is of particular importance in perennial species that will overwinter several times during their life-cycle. In temperate perennials prolonged exposure to cold regulates later stages of flowering such as uniform bud break in the spring, whereas in alpine species ensure floral formation and commitment before plants experience favorable environmental conditions for anthesis (Diggle, 1997; Lazaro *et al*., 2018; Meloche and Diggle, 2001). The maintenance of vegetative development after flowering, which is important for the perennial life strategy, is regulated by the seasonal cycling of floral repressors and the differential response of meristems to flower inductive stimuli due to age-related factors (Koskela *et al*., 2012; Wang *et al*., 2011; Wang *et al*., 2009). Here we characterized the role of the *A. alpina* floral repressor *PEP2*, the orthologue of the Arabidopsis *AP2*. Previous studies had demonstrated that *PEP2* regulates flowering through a *PEP1*–dependent and a *PEP1*–independent pathway (Bergonzi *et al*., 2013a). Our transcriptomic analysis indicated that *PEP2* influences the expression of genes involved in several developmental processes. Many of the identified genes though, might not be regulated directly by PEP2 but by complex downstream genetic interactions (Fig. 1 and Fig. S1). We also found both floral promoters and repressors differentially expressed in the *pep2* mutant. In Arabidopsis the AP2 protein was immunoprecipitated from the promoter region of *SOC1* (Yant *et al*., 2010). However, in our study the effect of *PEP2* on *AaSOC1* seems to be through *PEP1* (Fig. 1). To characterize the *PEP1*-dependent and the *PEP1*-independent role of *PEP2* on flowering we also employed physiological analysis and followed the expression of flowering time and meristem identity genes during the *A. alpina* life-cycle. These data indicated that *PEP2* regulates i) the age-dependent response to vernalization and ii) the temporal cycling of the floral repressor *PEP1* by ensuring the activation of *PEP1* expression after vernalization.

### PEP2 regulates the age-dependent response to vernalization

*PEP2* could rescue the early flowering phenotype of the Arabidopsis *ap2-7* mutant suggesting that its role on flowering time might be conserved (Fig. 2). In Arabidopsis, *AP2* is post-transcriptionally regulated by miR172, and *miR172b* is placed in the age pathway as it is transcriptionally controlled by the miR156 targets SPL9 and SPL15 (Hyun *et al*., 2016; Wu *et al*., 2009). AP2 also negatively regulates its own expression by directly binding to its own genomic locus, as well as to the loci of its regulators *miR156e*, *miR172b* and *FUL*, suggesting that *AP2* is transcriptionally regulated by multiple feedback loops (Balanza *et al*., 2018; Schwab *et al*., 2005; Yant *et al*., 2010). The AP2 protein was also immunoprecipitated from the chromatin of floral integrators and genes required for floral meristem development such as *SOC1*, *AGAMOUS* (*AG*) and *AP1* (Yant et al., 2010). The transcription of genes such as *SOC1* and *FUL* is also controlled by upstream regulators in the age pathway. SPL9 has been reported to bind to the first intron of *SOC1* and SPL15 to *FUL* and *miR172b* (Hyun et al., 2016; Wang et al., 2009). Overall, this complex genetic circuit that includes AP2 might contribute to the fast life-cycle of Arabidopsis, in which floral transition takes place soon after reproductive competence is acquired. Contrary to Arabidopsis, in *A. alpina* reproductive competence is uncoupled from flowering initiation. *A. alpina* plants become competent to flower after growing for five weeks in long day conditions but only initiate flowering when they are exposed to vernalization (Wang *et al*., 2009). This suggests that flowering in *A. alpina* is regulated by a strong interplay between the age and the vernalization pathways. Members of the SPL and AP2 families (e.g. *AaSPL15* and *AaTOE2*) are transcriptionally repressed by PEP1 in addition to the post-transcriptional and post-translational regulation by the microRNAs (Bergonzi *et al*., 2013a; Chen, 2004; Hyun *et al*., 2016; Mateos *et al*., 2017; Xu *et al*., 2016). Although, FLC in Arabidopsis targets a similar set of genes the strong interplay between the age and the vernalization pathway is most apparent in *A. alpina* (Deng *et al*., 2011; Mateos *et al*., 2017). Vernalization in *A. alpina* provides the condition where the age effect on flowering is apparent as it silences *PEP1*. Gradual changes in the accumulation of miR156 and the expression of the *SPL*s can be observed in the shoot apex of *A. alpina* plants that get older in LDs (Bergonzi *et al*., 2013a). However, the accumulation of downstream regulators in the age pathway, such as of the miR172, only increase in the shoot apex during vernalization and upon floral transition (Bergonzi *et al*., 2013a). Here we show that the expression of miR156 and of *AaSPL5* and *15* is not influenced in *pep2* plants grown in LDs (Fig. S3, Fig. 1E-G). Given that *PEP2* acts partially through *PEP1*, the lack of an effect in *pep2-1* on *AaSPL15* can be either due to the residual *PEP1* expression in the *pep2-1* mutant or to the existence of compensatory genetic mechanisms. Interestingly, *AaSPL9* mRNA levels were reduced in 6-week-old *pep2-1* seedlings compared to the wild type (Fig. S3). This effect of *PEP2* on *AaSPL9*, though, cannot be explained by the feedback loops described in Arabidopsis as *AaSPL9* transcript levels would be expected to be higher in *pep2-1* compared to the wild type (Fig. S3; Yant *et al*., 2010).

The *A. alpina* orthologue of *TFL1* (*AaTFL1*) has been previously reported to influence the effect of vernalization in an age-dependent manner, although its expression pattern does not differ between juvenile and adult apices before vernalization (Wang *et al*., 2011). Here we show that vernalization accelerated flowering in young *pep2-1* seedlings compared to *pep1-1*, suggesting *PEP2* also regulates the age-dependent response to vernalization in a *PEP1* independent pathways (Fig. 3). Interestingly, in our RNAseq analysis *AaTFL1* transcripts were reduced in the *pep2-1* mutant suggesting that *PEP2*, together with or through *AaTFL1*, sets an age threshold for flowering in response to vernalization. One major difference between *AaTFL1* and *PEP2*, though, is that lines with reduced *AaTFL1* activity do not flower without vernalization. These results suggest that *PEP2* plays additional roles in the regulation of flowering time in *A. alpina*. Transcriptomic experiments in Arabidopsis also showed that *TFL1* mRNA is down-regulated in *ap2* inflorescences compared to the wild type (Yant *et al*., 2010). However, no direct binding of AP2 to the *TFL1* locus has been detected by ChIP-Seq and therefore it is unclear whether there is a direct or indirect effect of AP2 on *TFL1* transcription (Yant *et al*., 2010).

### PEP2 ensures the activation of PEP1 after vernalization

Previous studies in *A. alpina* have demonstrated that *PEP2* regulates flowering in response to vernalization by enhancing the expression of *PEP1* (Bergonzi *et al*., 2013a). Here we show that the major role of *PEP2* in *PEP1* activation takes place after vernalization. *PEP1* expression in *A. alpina* is temporarily silenced during prolonged exposure to cold to define inflorescence fate, while it is up-regulated after vernalization to repress flowering in axillary branches and define the inflorescence fate (Lazaro *et al*., 2018; Wang *et al*., 2009). We have recently shown that the duration of vernalization influences *PEP1* reactivation in the shoot apex after the return to warm temperatures (Lazaro *et al*., 2018). Phenotypes correlated with high *PEP1* mRNA levels after vernalization (e.g. floral reversion and the presence of vegetative axillary branches) were compromised in the *pep2-1* mutant (Fig. 5; Lazaro et al., 2018). Accordingly, *PEP1* mRNA levels were reduced in vernalized *pep2-1* plants compared to the wild type both in the inflorescence stem and the axillary branches (Fig. 6; Wang et al., 2009; Lazaro et al., 2018). These results suggest that *PEP2* contributes to the perennial life-cycle and regulates perennial specific traits by activating *PEP1* after vernalization. In Arabidopsis the introgression of the *FRI* allele from the Sf-2 accession into Col extends the duration of cold temperatures required to silence *FLC* (Searle *et al*., 2006). Northern Arabidopsis accessions such as Lov-1 require several months of vernalization to achieve *FLC* silencing and similar to *A. alpina* Pajares, a shorter duration of cold temperatures causes *FLC* reactivation (Shindo *et al*., 2006). The link between *AP2* and *FLC* is not clear in Arabidopsis. AP2 does not bind to *FLC* locus in ChIP-seq experiments and in our study *FLC* expression was not altered in plants where the *ap2-7* mutant allele was introgressed into Col *FRI* Sf-2 background (Fig. S3; Yant et al., 2010). However, as the strongest difference in *PEP1* expression in the *pep2-1* mutant was after vernalization the effect of *AP2* in the Lov-1 accession should be analyzed to rule out a role of *AP2* on *FLC* reactivation after insufficient vernalization.

The unstable silencing of *FLC* involves changes in the accumulation of the H3 trimethylation at Lysine 27 (H3K27me3) (Angel *et al*., 2011; Coustham *et al*., 2012). The pattern of the H3K27me3 mark at the *PEP1* locus also correlates with changes in *PEP1* mRNA levels in *A. alpina* (Wang *et al*., 2009). *PEP1* shows a much higher and broader increase of H3K27me3 during the cold than *FLC*, and H3K27me3 levels rapidly decrease at *PEP1* after short vernalization periods (Angel *et al*., 2011; Lazaro *et al*., 2018; Wang *et al*., 2009). Although the proteins regulating histone modifications at the *PEP1* locus are not known, in Arabidopsis resetting of the epigenetic memory of *FLC* is dependent on the presence of TrxG components and the Jumonji C (JmjC) domain-containing demethylases EARLY FLOWERING 6 (ELF6) and RELATIVE OF EARLY FLOWERING 6 (REF6) (Crevillen *et al*., 2014; Noh *et al*., 2004; Yun *et al*., 2011). It has been shown that *AP2* has the ability to interact with a chromatin remodeling factor *HISTONE DEACETYLASE 19* (*HDA19*) to transcriptionally repress one of its targets (Krogan *et al*., 2012), but *AP2* has never been associated to histone demethylases.

We have recently demonstrated that *PEP1* is stably silenced in the shoot apical meristem of adult plants that commit to flowering during prolonged exposure to cold (Lazaro *et al*., 2018). In juvenile plants a similar length of vernalization fails to initiate flowering even if *PEP1* is silenced during cold (Lazaro *et al*., 2018). Floral commitment during vernalization is correlated with a higher expression of the floral meristem identity genes, *AaFUL, AaLFY* and *AaAP1* which are repressed by *PEP2* (Lazaro *et al*., 2018). This is evident by the precocious up-regulation of *AaFUL*, *AaLFY* and *AaAP1* mRNA levels in vernalized *pep2-1* plants compared to the wild type (Fig. 4 and S4). Although, the link between *PEP2* and *PEP1* resetting is not clear it seems that the achievement of floral commitment during vernalization is negatively correlated with *PEP1* up-regulation after the return to warm temperatures (Lazaro *et al*., 2018). In Arabidopsis, AP2 is not known to influence *FLC* transcription. However, *AP2* has been reported to be transcriptionally repressed by FUL and *FUL* overexpressing plants show reduced *FLC* expression (Balanz *et al*., 2014; Balanza *et al*., 2018). These results suggest that FUL might regulate *FLC* transcription independently or through *AP2*. These might also indicate that in *A. alpina* the role of PEP2 on *PEP1* expression might implicate other flowering time regulators, genes involved in the age pathway and genes ensuring floral commitment during vernalization. However, since PEP1 also transcriptionally regulates genes in these genetic pathways feedback mechanisms might also occur (Mateos *et al*., 2017).

## CONCLUSION

Our study demonstrates the instrumental role of *PEP2* in *A. alpina* regulating the age-dependent response to vernalization and facilitating the activation of *PEP1* after vernalization. As both roles of *PEP2* focus on whether floral commitment has been achieved during vernalization, they might not be completely independent. Upstream regulators of floral meristem identity genes such as *PEP2* might regulate the response to vernalization of individual meristems and contribute to the complex plant architecture of perennials.

## AKNOWLEDGEMENTS

We would like to thank SPP1530 and CEPLAS for funding to MCA, SPP1529 for funding to AP and KN and Purkyně fellowship from AS CR to AP. We would also like to thank Margaret Kox for critical reading of the manuscript.

## SUPPLEMENTARY DATA

**Dataset S1.** Transcripts identified as being differentially expressed in *pep2-1* compared to the wild type.

**Dataset S2.** Transcripts identified as being differentially expressed in *pep1-1* compared to the wild type.

**Dataset S3.** Transcripts identified as being differentially expressed in *pep2-1* compared to *pep1-1*.

**Table S1.** Primers used for PIPE-cloning of the *PEP2* locus

**Table S2.** Statistical differences in Figure S2 determined by multiple pairwise comparisons using Benjamini-Hochberg-corrected p-values comparing *FLC* mRNA levels between *FRI* and *FRI ap2-7* at different developmental stages.

**Table S3.** Statistical differences in Figure 6 determined by multiple pairwise comparisons using Benjamini-Hochberg-corrected p-values comparing *PEP1* mRNA levels between *pep2-1* and the wild type at different developmental stages.

**Fig. S1:** GO enriched categories in RNAseq experiment.

**Fig. S2.** *AP2* does not affect *FLC* expression in Arabidopsis.

**Fig. S3.** The expression level of *miR156*, *AaSPL5* and *AaSPL15* does not differ between wild type and *pep2-1* plants growing in long days.

**Fig. S4.** *PEP2* regulates the age-dependent response of *A. alpina* to vernalization.

**Fig. S5.** *PEP2* regulates *AaFUL*, *AaTFL1*, *AaLFY* and *AaAP1* expression during vernalization in adult plants.

---

**Fig. S1. GO enriched categories in RNAseq experiment.** Bubble network shows GO terms enriched among differentially expressed genes in *pep2-1*. Color represents p value, and size of the bubble represents the representation factor. Hypergeometric test Benjamini– Hochberg FDR correction. Cutoff 0.05. **(A)** Up regulated in *pep2-1* compared to the wild type (WT). **(B)** Up regulated in *pep2-1* compared to *pep1-1*. **(C)** Down regulated in *pep2-1* compared to the WT. **(D)** Down-regulated in *pep2-1* compared to *pep1-1*.

**Fig. S2. *AP2* does not affect *FLC* expression in Arabidopsis. (A)** Flowering time of *FRI* and *FRI ap2-7* plants scored as the number of leaves at flowering. Dark grey color represents rosette leaves and light grey color cauline leaves. The asterisk stands for a significant difference in the total leaf number determined by a Student T-test (p-value<0.01). **(B)** Relative expression of *FLC* in the shoot apical meristem of *FRI* and *FRI ap2-7* plants before, during and after 40 days of vernalization. There are no significant differences between *FRI* and *FRI ap2-7* samples at each time point determined by multiple pairwise comparisons using Benjamini-Hochberg-corrected p-values (α-value of 0.05). Detailed information on significant differences can be found in Table S2. Error bars indicate s.d.m.

**Fig. S3. The expression level of *miR156*, *AaSPL5* and *AaSPL15* does not differ between wild type and *pep2-1* plants growing in long days.** Relative expression of miR156 **(A)**, *AaSPL5* **(B)**, *AaSPL9* **(C)** and *AaSPL15* **(D)** in wild type (WT) and the *pep2-1* mutant. Apices were harvested from WT and *pep2-1* seedlings growing for 3, 4 and 6 weeks in LDs. Asterisks stand for significant differences between WT and *pep2-1* at each time point determined by multiple pairwise comparisons using Benjamini-Hochberg-corrected p-values (α-value of 0.05). Values are the average of 2 biological replicates, error bars indicate s.d.m.

**Fig. S4. *PEP2* regulates the age-dependent response of *A. alpina* to vernalization.** Flowering time demonstrated as the number of days to flower emergence of 3-week-old wild type (WT), *pep1-1* mutant and *pep2-1* mutant, vernalized for 12 weeks. WT did not flower (NF). Error bars indicate s.d.m.

**Fig. S5. *PEP2* regulates *AaFUL*, *AaTFL1*, *AaLFY* and *AaAP1* expression during vernalization in adult plants.** Relative expression of *AaFUL* **(A)**, *AaLFY* **(B)**, *AaAP1* **(C).** 6-week-old wild type (WT) and *pep2-1* shoot apices were harvested before and during 12 weeks of vernalization. Asterisks stand for significant differences between the WT and *pep2-1* at each time point determined by multiple pairwise comparisons using Benjamini-Hochberg-corrected p-values (α-value of 0.05). Error bars indicate s.d.m.

**Table S1.** Primers used for PIPE-cloning of the *PEP2* locus

**Table S2.** Statistical differences in Figure S2 determined by multiple pairwise comparisons using Benjamini-Hochberg-corrected p-values comparing *FLC* mRNA levels between *FRI* and *FRI ap2-7* at different developmental stages. The comparisons highlighted in yellow are significantly different (α-value of 0.05).

**Table S3.** Statistical differences in Figure 6 determined by multiple pairwise comparisons using Benjamini-Hochberg-corrected p-values comparing *PEP1* mRNA levels between *pep2-1* and the wild type at different developmental stages. The comparisons highlighted in yellow are significantly different (α-value of 0.05).

## ABBREVIATIONS

DAG: Days after germination
LDs: Long days
SDs: Short days

## REFERENCES

Amasino R. 2009. Floral induction and monocarpic versus polycarpic life histories. Genome Biology 10.

Angel A, Song J, Dean C, Howard M. 2011. A Polycomb-based switch underlying quantitative epigenetic memory. Nature 476, 105–108.

Aukerman MJ, Sakai H. 2003. Regulation of flowering time and floral organ identity by a microRNA and its *APETALA2*-like target genes. Plant Cell 15, 2730–2741.

Balanz V, Martnez-Fernndez I, Ferrndiz C. 2014. Sequential action of *FRUITFULL* as a modulator of the activity of the floral regulators *SVP* and *SOC1*. Journal of Experimental Botany 65, 1193–1203.

Balanza V, Martinez-Fernandez I, Sato S, Yanofsky MF, Kaufmann K, Angenent GC, Bemer M, Ferrandiz C. 2018. Genetic control of meristem arrest and life span in Arabidopsis by a FRUITFULL-APETALA2 pathway. Nature Communications 9.

Benjamini Y, Hochberg Y. 1995. Controlling the False Discovery Rate - a Practical and Powerful Approach to Multiple Testing. Journal of the Royal Statistical Society Series B-Methodological 57, 289–300.

Bergonzi S, Albani MC. 2011. Reproductive competence from an annual and a perennial perspective. Journal of Experimental Botany 62, 4415–4422.

Bergonzi S, Albani MC, van Themaat EVL, Nordstrom KJV, Wang RH, Schneeberger K, Moerland PD, Coupland G. 2013. Mechanisms of Age-Dependent Response to Winter Temperature in Perennial Flowering of *Arabis alpina*. Science 340, 1094–1097.

Billings WD, Mooney HA. 1968. Ecology of Arctic and Alpine Plants. Biological Reviews of the Cambridge Philosophical Society 43, 481–529.

Bowman JL, Smyth DR, Meyerowitz EM. 1991. Genetic Interactions among Floral Homeotic Genes of Arabidopsis. Development 112, 1–20.

Castro PH, Couto D, Freitas S, Verde N, Macho A, Huguet S, Botella MA, Ruiz-Albert J, Tavares RM, Bejarano ER, Azevedo H. 2016. SUMO proteases ULP1c and ULP1d are required for development and osmotic stress responses in *Arabidopsis thaliana*. Plant Molecular Biology 92, 143–159.

Chen XM. 2004. A microRNA as a translational repressor of *APETALA2* in *Arabidopsis* flower development. Science 303, 2022–2025.

Cline MS, Smoot M, Cerami E, Kuchinsky A, Landys N, Workman C, Christmas R, Avila-Campilo I, Creech M, Gross B, Hanspers K, Isserlin R, Kelley R, Killcoyne S, Lotia S, Maere S, Morris J, Ono K, Pavlovic V, Pico AR, Vailaya A, Wang PL, Adler A, Conklin BR, Hood L, Kuiper M, Sander C, Schmulevich I, Schwikowski B, Warner GJ, Ideker T, Bader GD. 2007. Integration of biological networks and gene expression data using Cytoscape. Nature Protocols 2, 2366–2382.

Clough SJ, Bent AF. 1998. Floral dip: a simplified method for Agrobacterium-mediated transformation of *Arabidopsis thaliana*. Plant Journal 16, 735–743.

Conti L, Price G, O’Donnell E, Schwessinger B, Dominy P, Sadanandom A. 2008. Small Ubiquitin-Like Modifier Proteases OVERLY TOLERANT TO SALT1 and-2 Regulate Salt Stress Responses in *Arabidopsis*. Plant Cell 20, 2894–2908.

Coustham V, Li PJ, Strange A, Lister C, Song J, Dean C. 2012. Quantitative Modulation of Polycomb Silencing Underlies Natural Variation in Vernalization. Science 337, 584–587.

Crevillen P, Sonmez C, Wu Z, Dean C. 2013. A gene loop containing the floral repressor *FLC* is disrupted in the early phase of vernalization. Embo Journal 32, 140–148.

Crevillen P, Yang HC, Cui X, Greeff C, Trick M, Qiu Q, Cao XF, Dean C. 2014. Epigenetic reprogramming that prevents transgenerational inheritance of the vernalized state. Nature 515, 587–590.

Czechowski T, Stitt M, Altmann T, Udvardi MK, Scheible WR. 2005. Genome-wide identification and testing of superior reference genes for transcript normalization in Arabidopsis. Plant Physiology 139, 5–17.

Deng WW, Ying H, Helliwell CA, Taylor JM, Peacock WJ, Dennis ES. 2011. FLOWERING LOCUS C (FLC) regulates development pathways throughout the life cycle of *Arabidopsis*. Proceedings of the National Academy of Sciences of the United States of America 108, 6680–6685.

Diggle PK. 1997. Extreme preformation in alpine *Polygonum viviparum*: An architectural and developmental analysis. American Journal of Botany 84, 154–169.

Guo D, Wong WS, Xu WZ, Sun FF, Qing DJ, Li N. 2011. *Cis-cinnamic acid-enhanced 1* gene plays a role in regulation of Arabidopsis bolting. Plant Molecular Biology 75, 481–495.

Hyun Y, Richter R, Vincent C, Martinez-Gallegos R, Porri A, Coupland G. 2016. Multi-layered Regulation of SPL15 and Cooperation with SOC1 Integrate Endogenous Flowering Pathways at the *Arabidopsis* Shoot Meristem. Developmental Cell 37, 254–266.

Keith RA, Mitchell-Olds T. 2017. Testing the optimal defense hypothesis in nature: Variation for glucosinolate profiles within plants. Plos One 12, e0180971.

Klock HE, Koesema EJ, Knuth MW, Lesley SA. 2008. Combining the polymerase incomplete primer extension method for cloning and mutagenesis with microscreening to accelerate structural genomics efforts. Proteins-Structure Function and Bioinformatics 71, 982–994.

Koskela EA, Mouhu K, Albani MC, Kurokura T, Rantanen M, Sargent DJ, Battey NH, Coupland G, Elomaa P, Hytonen T. 2012. Mutation in *TERMINAL FLOWER1* Reverses the Photoperiodic Requirement for Flowering in the Wild Strawberry *Fragaria vesca*. Plant Physiology 159, 1043–1054.

Kotoda N, Iwanami H, Takahashi S, Abe K. 2006. Antisense expression of *MdTFL1*, a *TFL1*-like gene, reduces the juvenile phase in apple. Journal of the American Society for Horticultural Science 131, 74–81.

Krogan NT, Hogan K, Long JA. 2012. APETALA2 negatively regulates multiple floral organ identity genes in *Arabidopsis* by recruiting the co-repressor TOPLESS and the histone deacetylase HDA19. Development 139, 4180–4190.

Lazaro A, Obeng-Hinneh E, Albani MC. 2018. Extended Vernalization Regulates Inflorescence Fate in *Arabis alpina* by Stably Silencing *PERPETUAL FLOWERING1*. Plant Physiology 176, 2819–2833.

Lee I, Amasino RM. 1995. Effect of Vernalization, Photoperiod, and Light Quality on the Flowering Phenotype of Arabidopsis Plants Containing the *FRIGIDA* Gene. Plant Physiology 108, 157–162.

Maere S, Heymans K, Kuiper M. 2005. BiNGO: a Cytoscape plugin to assess overrepresentation of Gene Ontology categories in Biological Networks. Bioinformatics 21, 3448–3449.

Mateos JL, Tilmes V, Madrigal P, Severing E, Richter R, Rijkenberg CWM, Krajewski P, Coupland G. 2017. Divergence of regulatory networks governed by the orthologous transcription factors FLC and PEP1 in Brassicaceae species. Proceedings of the National Academy of Sciences of the United States of America 114, E11037–E11046.

Mathieu J, Yant LJ, Murdter F, Kuttner F, Schmid M. 2009. Repression of Flowering by the miR172 Target SMZ. Plos Biology 7, e1000148.

Meloche CG, Diggle PK. 2001. Preformation, architectural complexity, and developmental flexibility in *Acomastylis rossii* (Rosaceae). American Journal of Botany 88, 980–991.

Michaels SD, Amasino RM. 1999. *FLOWERING LOCUS C* encodes a novel MADS domain protein that acts as a repressor of flowering. Plant Cell 11, 949–956.

Mohamed R, Wang CT, Ma C, Shevchenko O, Dye SJ, Puzey JR, Etherington E, Sheng XY, Meilan R, Strauss SH, Brunner AM. 2010. *Populus CEN/TFL1* regulates first onset of flowering, axillary meristem identity and dormancy release in *Populus*. Plant Journal 62, 674–688.

Noh B, Lee SH, Kim HJ, Yi G, Shin EA, Lee M, Jung KJ, Doyle MR, Amasino RM, Noh YS. 2004. Divergent roles of a pair of homologous jumonji/zinc-finger-class transcription factor proteins in the regulation of Arabidopsis flowering time. Plant Cell 16, 2601–2613.

Nördstrom KJ, Albani MC, James GV, Gutjahr C, Hartwig B, Turck F, Paszkowski U, Coupland G, Schneeberger K. 2013. Mutation identification by direct comparison of whole-genome sequencing data from mutant and wild-type individuals using *k*-mers. Nat Biotechnol 31, 325–330.

Nordstrom KJV, Albani MC, James GV, Gutjahr C, Hartwig B, Turck F, Paszkowski U, Coupland G, Schneeberger K. 2013. Mutation identification by direct comparison of whole-genome sequencing data from mutant and wild-type individuals using k-mers. Nature Biotechnology 31, 325–330.

Park JY, Kim H, Lee I. 2017. Comparative analysis of molecular and physiological traits between perennial *Arabis alpina* Pajares and annual *Arabidopsis thaliana* Sy-0. Scientific Reports 7, 13348.

Schmid M, Uhlenhaut NH, Godard F, Demar M, Bressan R, Weigel D, Lohmann JU. 2003. Dissection of floral induction pathways using global expression analysis. Development 130, 6001–6012.

Schwab R, Palatnik JF, Riester M, Schommer C, Schmid M, Weigel D. 2005. Specific effects of MicroRNAs on the plant transcriptome. Developmental Cell 8, 517–527.

Searle I, He YH, Turck F, Vincent C, Fornara F, Krober S, Amasino RA, Coupland G. 2006. The transcription factor FLC confers a flowering response to vernalization by repressing meristem competence and systemic signaling in *Arabidopsis*. Genes & Development 20, 898–912.

Sheldon CC, Rouse DT, Finnegan EJ, Peacock WJ, Dennis ES. 2000. The molecular basis of vernalization: The central role of *FLOWERING LOCUS C (FLC)*. Proceedings of the National Academy of Sciences of the United States of America 97, 3753–3758.

Shindo C, Lister C, Crevillen P, Nordborg M, Dean C. 2006. Variation in the epigenetic silencing of *FLC* contributes to natural variation in Arabidopsis vernalization response. Genes & Development 20, 3079–3083.

Trapnell C, Pachter L, Salzberg SL. 2009. TopHat: discovering splice junctions with RNA-Seq. Bioinformatics 25, 1105–1111.

Trapnell C, Williams BA, Pertea G, Mortazavi A, Kwan G, van Baren MJ, Salzberg SL, Wold BJ, Pachter L. 2010. Transcript assembly and quantification by RNA-Seq reveals unannotated transcripts and isoform switching during cell differentiation. Nature Biotechnology 28, 511–U174.

Wang RH, Albani MC, Vincent C, Bergonzi S, Luan M, Bai Y, Kiefer C, Castillo R, Coupland G. 2011. *Aa TFL1* Confers an Age-Dependent Response to Vernalization in Perennial *Arabis alpina*. Plant Cell 23, 1307–1321.

Wang RH, Farrona S, Vincent C, Joecker A, Schoof H, Turck F, Alonso-Blanco C, Coupland G, Albani MC. 2009. *PEP1* regulates perennial flowering in *Arabis alpina*. Nature 459, 423–427.

Willing EM, Rawat V, Maumus F, James GV, Nordström KJV, Becker C, Warthmann N, Chica C, Szarzynska B, Zytnicki M, Albani MC, Kiefer C, Bergonzi S, Castaings L, Mateos JL, Berns MC, Bujdoso N, Piofczyk T, de Lorenzo L, Barrero-Sicilia C, Mateos I, Piednoël M, Hagmann J, Chen-Min-Tao R, Iglesias-Fernández R, Schuster SC, Alonso-Blanco C, Roudier F, Carbonero P, Paz-Ares J, Davis SJ, Pecinka A, Quesneville H, Colot V, Lysak MA, Weigel D, Coupland G, Schneeberger K. 2015. Lack of symmetric CG methylation and long-lasting retrotransposon activity have shaped the genome of *Arabis alpina*, Nature Plants 1, 14023.

Wu G, Park MY, Conway SR, Wang JW, Weigel D, Poethig RS. 2009. The sequential action of miR156 and miR172 regulates developmental timing in Arabidopsis. Cell 138, 750–759.

Wu G, Poethig RS. 2006. Temporal regulation of shoot development in *Arabidopsis thaliana* by *miR156* and its target *SPL3*. Development 133, 3539–3547.

Xu ML, Hu TQ, Zhao JF, Park MY, Earley KW, Wu G, Yang L, Poethig RS. 2016. Developmental Functions of miR156-Regulated *SQUAMOSA PROMOTER BINDING PROTEIN-LIKE (SPL)* Genes in *Arabidopsis thaliana*. Plos Genetics 12, e1006263.

Yant L, Mathieu J, Dinh TT, Ott F, Lanz C, Wollmann H, Chen XM, Schmid M. 2010. Orchestration of the Floral Transition and Floral Development in Arabidopsis by the Bifunctional Transcription Factor APETALA2. Plant Cell 22, 2156–2170.

Yun H, Hyun Y, Kang MJ, Noh YS, Noh B, Choi Y. 2011. Identification of regulators required for the reactivation of *FLOWERING LOCUS C* during Arabidopsis reproduction. Planta 234, 1237–1250.

